# Invertebrate herbivory damage of lowland plant species decreases after an experimental shift to higher altitudes

**DOI:** 10.1101/2023.02.27.530180

**Authors:** Karolína Hrubá, Dagmar Hucková, Jan Klečka

## Abstract

Many species of plants and animals shift to higher altitudes in response to the ongoing climate warming. Such shifts of species distributions lead to the co-occurrence of species that have not previously lived in the same environment and allow the emergence of novel plant-animal interactions with potential implications for species diversity and community composition in mountain habitats. According to the enemy release hypothesis, the spread of plants in new geographic regions may be facilitated by the reduction of damage caused by natural enemies, such as herbivores. While the importance of this mechanism for the spread of invasive exotic species has been established, it is unclear whether the movement of plants uphill within their native region in response to increasing temperatures may be also facilitated by the reduction of herbivory at sites above their current upper altitudinal limit. In our study, we experimentally tested this hypothesis. We compared herbivory damage of six species of lowland plants grown in pots exposed to herbivores at their native sites in the lowland and at sites above their current upper altitudinal limit. As a control, we also measured herbivory damage of six plants growing naturally across the entire range of altitude. We found that lowland plants had reduced herbivory damage when they were moved to highland sites, while herbivory damage of species naturally growing at both altitudes did not differ. Changes of herbivory damage were modulated by leaf dry matter content and to a lesser degree also by specific leaf area and plant height. Our results provide support for the enemy release hypothesis in the novel context of altitudinal range shifts. We conclude that the reduction of herbivory damage may facilitate the spread of plants above their current upper altitudinal limit in response to increasing temperature.

## Introduction

Global warming leads to large-scale changes of the geographic distribution of many organisms, such as shifts to higher altitudes or towards the poles (Kerr et al. 2015; Freeman et al. 2018; Fazlioglu et al. 2020). Such range shifts lead to the emergence of communities with novel species combinations with important consequences for local biodiversity (Hobbs et al. 2006; Hobbs et al. 2009). For example, the spread of herbivores to higher altitudes can affect the composition of mountain plant communities (Descombes et al. 2020). The spread of plants uphill can affect mountain species which have to compete for resources (Nomoto & Alexander 2021) or for pollinators (Hernández-Castellano et al. 2020) with the incoming plants. However, our knowledge of these processes and their long-term ecological impacts is very limited and there is a number of open questions related to factors affecting the establishment and spread of new species in mountain habitats and their direct and indirect effects on mountain communities mediated by interactions with species at other trophic levels (Tomiolo & Ward 2018; Nomoto & Alexander 2021).

Causes and ecological consequences of the spread of plants to new geographic regions were studied mostly in invasive exotic species (Vilà et al. 2009; Simberloff et al. 2013). According to the enemy release hypothesis, their spread to new areas can be facilitated by the reduction of damage caused by herbivores or pathogens because the plants escape specialized natural enemies occurring in their native range (Kambo & Kotanen 2014; Keane & Crawley 2002; Costan et al., 2022). Herbivores attacking the exotic plants in their invasive range are usually generalists which may not be adapted to cope with chemical defences of the exotic plants (Cappuccino & Carpenter 2005; Enge et al. 2012). Lower aboveground or belowground herbivory damage provides a competitive advantage to exotic plants, which facilitates their spread (Keane & Crawley 2002; Engelkes et al 2008). Such reduction of herbivory was detected also at smaller scales, e.g., in populations of invasive species at the margins of their geographic range during range expansion at the intra-continental scale (Morriën et al. 2010; Kambo & Kotanen 2014). We hypothesize that a similar reduction of herbivory may facilitate the spread of plants to higher altitudes within their native range in response to climate change.

Altitudinal range shifts are a striking case when the reorganization of local community structure can be observed at a small spatial scale because of the steep temperature gradient in the mountains. The responses of individual species are highly variable (Parmesan & Yohe 2003). While the optimum altitude for some plant species recorded by Lenoir et al. (2008) shifted by several hundred meters over a few decades, it changed little or even decreased in other cases. Hence, altitudinal range shifts do not represent the movement of entire communities upwards. Rather, species move at a different pace, which creates novel combinations of species that have not coexisted previously in high-altitude habitats. This trend has been accelerating during the last few decades, so mountain communities are increasingly affected by an influx of novel species from lower altitudes (Walther et al. 2005; Steinbauer et al. 2018). However, we do not know well how such shifts affect the interactions of the plants with herbivores, pollinators, or seed dispersers (Tomiolo & Ward 2018; Hernández-Castellano et al. 2020).

Native species, originating in lowland communities, shifting to higher altitudes move over a relatively short distance, but still represent novel species in the high-altitude communities. The main difference compared to invasive exotic species, which come from distant regions, is that mountain communities could contain species that are phylogenetically closely related to plants coming from the lowland. This raises the question whether the shifts of lowland plants to higher altitudes may be facilitated by reduced herbivory as observed in invasive exotic species (Kambo & Kotanen 2014; Keane & Crawley 2002; Costan et al. 2022). In addition, the abundance and activity of generalist herbivorous invertebrates often decreases with increasing elevation because of less favourable climate compared to the lowlands (Scheidel & Bruelheide 2001; Scheidel et al. 2003; Moreira et al. 2017), which may result in lower herbivory damage at higher elevations (Reynolds & Crossley 1997; Scheidel et al. 2003; Pellisier et al. 2014; Galmán et al. 2018).

Herbivory damage can be affected not only by phylogenetic relatedness and increasing elevation but also by plant functional traits (Pérez-Harguindeguy et al. 2003). Higher leaf dry matter content (LDMC) typical for plants whose leaves are tougher, have higher longevity, and slow growth decreases leaf palatability for herbivores. On the contrary, leaves with high specific leaf area (SLA), which are thinner and softer, are often preferred by herbivores (Reese et al. 2016). Also, the height of a plant is an important indicator of invertebrate herbivory, because some flightless herbivores, in particular slugs and snails, forage close to the ground (Hahn et al. 2011; Hrubá et al. 2022). On the other hand, taller plants can be more visible and thus attractive to flying herbivores (Feeny 1976). In general, plant species with tough leaves with high LDMC, low SLA, and taller size are less attractive for herbivores and generally have lower herbivory damage (Reese et al. 2016; Těšitel et al. 2021; Hrubá et al. 2022).

Despite the ubiquitous nature of altitudinal range shifts and their potential impact on plant-animal interactions, we are not aware of any empirical studies focusing on the impact of novel species on the relationships of plants with herbivores, with the exception of a study by Descombes et al. (2020) who transferred herbivorous locusts to higher altitudes. Therefore, we conducted an experimental study where we transferred six species of lowland thermophilous plants to altitudes above their current upper altitudinal limit and tested whether this led to a reduction of their herbivory damage using plants naturally growing across the entire range of altitude as a control. We also aimed to test the effect of several plant traits known to affect herbivory, such as leaf dry matter content (LDMC), specific leaf area (SLA), and plant height (Descombes et al. 2017; Těšitel et al. 2021; Wang et al. 2022). Specifically, we asked the following questions:

1. Do lowland species suffer lower damage by herbivores when experimentally moved to altitudes above their current upper altitudinal limit, as predicted by the enemy release hypothesis?
2. Are plant species naturally growing along the entire range of altitude more damaged by herbivores in the lowland where we expect higher abundance of herbivores?
3. Which plant traits influence herbivory damage and its differences between the lowland and mountain sites?

## Material and methods

### Study sites

The study was carried out in the southern part of the Czech Republic. We conducted our experiment at 14 lowland sites (528 – 609 m a.s.l.) in the surroundings of the city of Český Krumlov and six highland sites (856 – 1020 m a.s.l.) in the nearby Šumava Mountains (Table 1, Figure 1). In both cases, the sites were meadows with a relatively high diversity of plant species. The choice of lowland sites was based on the occurrence of plant species selected for the experiment (six widely distributed species, see below). We aimed to include six sites where each plant species grows, although some species were restricted to a smaller number of sites. Each lowland site contained one to four target species growing there naturally, none of the sites contained all six selected plant species. Therefore, we had to sample more sites in the lowland in comparison to the highland. The highland sites were all located at an altitude higher than the highest known occurrence of the target lowland species in the region based on the Pladias database (Chytrý et al. 2021). Several sites were located on public land inside protected landscape areas with a low level of protection (Blanský Les and Šumava), so conducting the experiments was possible with the permission of their management. The remaining sites were on private land, and we obtained permission for our work from the owners.

**Table 1.** The list of study sites in the lowland and the highland. Altitude and GPS coordinates of the centre of each site are provided.

| Site ID | Site name | Altitude (m a.s.l.) | GPS |
| --- | --- | --- | --- |
| Highland 1 | Arnoštov | 856 | 48.89307N, 13.99915E |
| Highland 2 | Horní Sněžná | 1020 | 48.88743N, 13.93954E |
| Highland 3 | Čtyři domy | 956 | 48.89779N, 13.95513E |
| Highland 4 | Strážný 1 | 982 | 48.93432N, 13.71026E |
| Highland 5 | Strážný 2 | 973 | 48.92935N, 13.73084E |
| Highland 6 | Strážný 3 | 910 | 48.92762N, 13.71185E |
| Lowland 1 | Brloh | 588 | 48.92492N, 14.22733E |
| Lowland 2 | Cvičák | 570 | 48.82559N, 14.31408E |
| Lowland 3 | Kájov | 553 | 48.81425N, 14.27224E |
| Lowland 4 | Kalamandra | 543 | 48.82179N, 14.27911E |
| Lowland 5 | Křenovský dvůr | 566 | 48.82147N, 14.24758E |
| Lowland 6 | Lazec | 609 | 48.83522N, 14.26014E |
| Lowland 7 | Liščí hora | 573 | 48.82858N, 14.32326E |
| Lowland 8 | Český Krumlov 1 | 543 | 48.82323N, 14.32137E |
| Lowland 9 | Český Krumlov 2 | 528 | 48.81838N, 14.30662E |
| Lowland 10 | Český Krumlov 3 | 530 | 48.81854N, 14.30744E |
| Lowland 11 | Vyšný 1 | 579 | 48.82637N, 14.30220E |
| Lowland 12 | Vyšný 2 | 581 | 48.83102N, 14.29278E |
| Lowland 13 | Vyšný 3 | 590 | 48.83088N, 14.30778E |
| Lowland 14 | Vyšný 4 | 578 | 48.83046N, 14.29691E |

**Figure 1.**
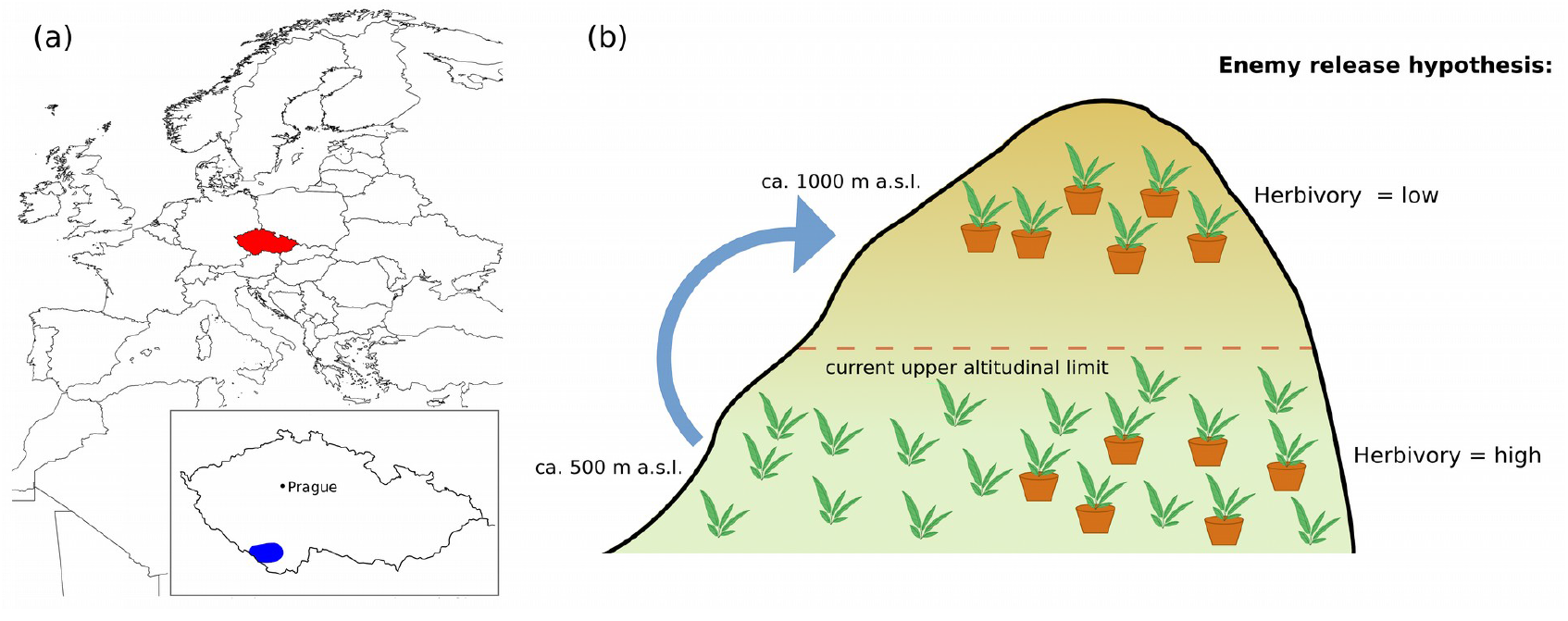
The location of the study area and the scheme of our experimental design. (a) The study sites were located in the southern part of the Czech Republic in the area shown in blue. The map was plotted using the rworldmap package in R (South, 2011). Map lines delineate study areas and do not necessarily depict accepted national boundaries. (b) Thermophilous lowland plants were exposed in pots at lowland sites where they grow naturally and at highland sites above their current upper altitudinal limit.

### Study species

We selected two groups of plant species (Table 2) for our study. The first group was experimental, grown in pots. This group consists of six thermophilous species from three families (Asteraceae: *Centaurea jacea, C. scabiosa*, and *Senecio jacobea*; Lamiaceae: *Origanum vulgare and Salvia verticillata*; and Plantaginaceae: *Veronica teucrium*) naturally growing only at the lowland sites. The upper altitudinal limit of these species in the region is lower than the altitude of the highland sites chosen for the experiment (Chytrý et al. 2021). We collected seeds of this group of experimental plant species at the lowland sites in August – September 2020. We sowed the seeds into germination trays with a mixture of garden soil, compost, and sand (2:1:1 ratio) with the addition of a small quantity of limestone in March 2021. We transferred the seedlings individually into pots (diameter 11 cm) about two weeks after germination and grew them in a greenhouse to prevent access by herbivores.

**Table 2.** The list of plant species used and the summary of functional trait data. Origin distinguishes potted plants growing originally in the lowland (Potted) and plants naturally growing at both altitudes (Natural). Mean value of three functional traits measured in the end of the experiment is also provided for each species; separately for the lowland and highland sites. LDMC is expressed as the proportion of dry matter in plant leaves, i.e., dry weight (g) / fresh weight (g). SLA is expressed as the proportion of leaf area (mm^2^) / leaf mass (mg).

| Species | Origin | Lowland |  |  | Highland |  |  |
| --- | --- | --- | --- | --- | --- | --- | --- |
|  |  | Height (cm) | SLA (mm<sup>2</sup>/mg) | LDMC | Height (cm) | SLA (mm<sup>2</sup>/mg) | LDMC |
| *Fragaria* spp. | Natural | 17.78 | 20.14 | 0.3858 | 17.18 | 29.25 | 0.3284 |
| *Campanula patula* L. | Natural | 29.43 | 36.85 | 0.2095 | 35.69 | 40.38 | 0.1778 |
| *Hypericum* spp. | Natural | 39.94 | 26.26 | 0.3166 | 32.63 | 30.87 | 0.3118 |
| *Plantago lanceolata* L. | Natural | 23.05 | 14.12 | 0.1959 | 23.52 | 18.77 | 0.1843 |
| *Veronica chamaedris* L. | Natural | 18.92 | 30.57 | 0.2484 | 21.04 | 27.55 | 0.2837 |
| *Knautia arvensis* (L.) Coult. | Natural | 48.14 | 18.47 | 0.2062 | 60.07 | 18.35 | 0.2011 |
| *Centaurea jacea* L. | Potted | 10.14 | 22.72 | 0.1472 | 14.02 | 23.84 | 0.1335 |
| *Centaurea scabiosa* L. | Potted | 12.83 | 22.07 | 0.1330 | 15.33 | 23.10 | 0.1252 |
| *Salvia verticillata* L. | Potted | 9.40 | 26.83 | 0.1375 | 15.24 | 33.03 | 0.1209 |
| *Veronica teucrium* L. | Potted | 9.45 | 37.07 | 0.1723 | 7.55 | 34.94 | 0.1786 |
| *Origanum vulgare* L. | Potted | 16.35 | 31.17 | 0.2251 | 16.60 | 35.64 | 0.2004 |
| *Senecio jacobea* L. | Potted | 6.51 | 24.51 | 0.1201 | 9.54 | 29.23 | 0.1045 |

The second group of plant species, which served as a control in the experiment, consisted of six plant taxa which grow naturally at the lowland as well as the highland sites. We are aware of the limitations of using these naturally growing species as a control. This group of species should be ideally grown under the same conditions and planted in the pots in the same way as the first group. However, it was not feasible due to logistical constraints. These species from five families (Rosaceae: *Fragaria* spp.; Campanulaceae: *Campanula patula*; Plantaginaceae: *Veronica chamaedrys and Plantago lanceolata*; Dipsacaceae: *Knautia arvensis*; and Hypericaceae: *Hypericum* spp.) (Table 2) were selected on the basis of their high abundance at our focal lowland and highland sites. Each of them occurred at multiple lowland sites (two to seven, mean 4.5 sites) and highland sites (two to six, mean 4.5 sites). The selection included four taxa identified at the species level and two identified at the genus level. Specifically, *Fragaria* spp. and *Hypericum* spp. include the mixture of two very similar species (*Fragaria vesca* and *F. viridis*; *Hypericum perforatum* and *H. maculatum*), which we consider equivalent for the purpose of our analyses. The presence of the same species or genera at both lowland and highland sites allowed us to compare the natural level of herbivory between the two altitudes.

### Experimental design and data collections

We conducted the field experiment during June–July 2021. We placed 15 individuals of each potted species at six lowland sites (reduced to five sites in two of the species) where the species grows naturally and also at all six highland sites. The pots were distributed in the meadow in three loose clusters per one species set >5 m apart, each containing five pots arranged in a 1x1 m square. Each pot was secured by a wire stuck into the ground, which prevented the fall of the pot, e.g., in the case of strong wind. We left the pots with seedlings exposed to herbivores at experimental sites for exactly three weeks.

After three weeks, we measured the height of all potted individuals. Then, we cut the whole individuals and placed them individually in zip lock plastic bags with a piece of filter paper soaked with water. We also randomly chose 15 individuals of the selected plant species naturally growing at both lowland and highland sites (Table 2), measured their height, cut them at ground level, and also put them in zip lock bags. The bags with the plants were stored in polystyrene boxes with ice preventing the wilting of plants and enabling the leaves to fully saturate by water. In the laboratory, we cut all leaves of each plant individual. The leaves were weighed immediately after releasing from zip lock plastic bags to obtain fresh weight of each individual, followed by scanning of the leaves within one day. Afterwards, all scanned leaves were dried in the oven at 70°C for 72 hours (Pérez-Harguindeguy et al. 2003) and dry weight was then measured. We used ImageJ software to measure the total area of the scanned leaf for each individual. If present, the parts of the scanned leaves that appeared to be eaten by herbivores were manually completed to resemble the original shape of the leaves. The corresponding area was calculated using the ImageJ software. Herbivory damage was expressed as 1 – (actual leaf area / leaf area of the completed leaf), i.e., the proportion of the leaf area missing, apparently due to herbivory (Hrubá et al. 2022). We also measured specific leaf area (SLA), and leaf dry matter content (LDMC) following the protocol of Pérez-Harguindeguy et al. (2003).

### Data analysis

We conducted all analyses in the R software version 4.0.3 (R Core Team 2020) using linear mixed effect models (LMM) using *lme4* (Bates et al. 2015) and *lmerTest* (Kuznetsova et al. 2017) packages. We tested the effect of plant origin (naturally growing vs. potted plants originating in the lowland), altitude (lowland vs. highland), and plant traits on herbivory damage, including interactions shown in Table 3. Herbivory damage, the response variable, was expressed as the proportion of the leaf area eaten by herbivores and was logit-transformed prior to analysis. The explanatory variables plant height and SLA were log-transformed and LDMC, expressed as the proportion of leaf dry weight / fresh weight, was logit-transformed. Plant species, site, and cluster nested within site were used as random effects.

**Table 3.**
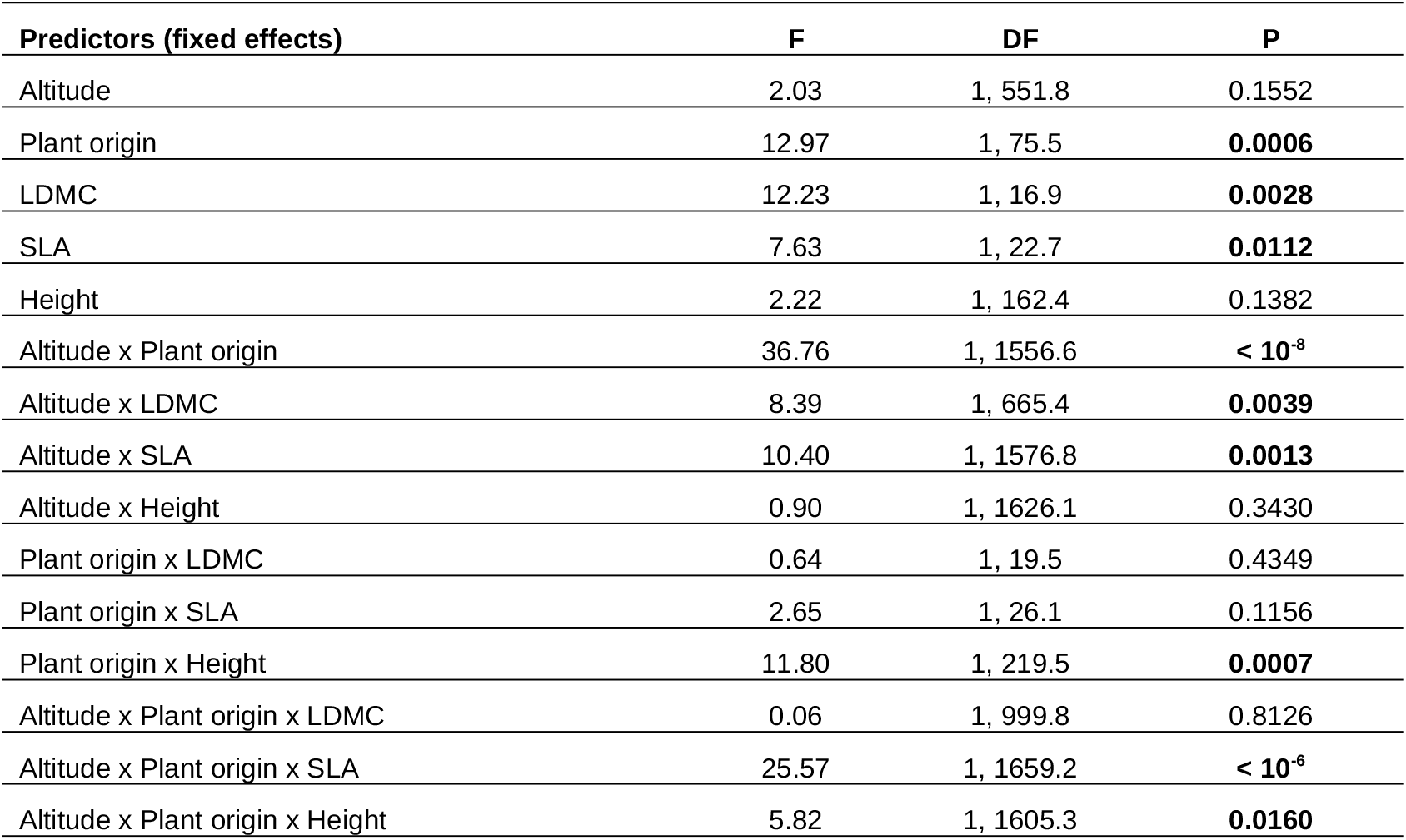
The results of the linear mixed effects model testing the dependence of the herbivory damage on plant origin, altitude, and plant traits. The model also included species, site, and cluster nested within site as random effects. Herbivory damage, the response variable, was logit-transformed prior to analysis. The explanatory variables plant height and SLA were log-transformed and LDMC was logit-transformed. Model estimates are plotted in Fig. 2. DF = degrees of freedom.

| Predictors (fixed effects) | F | DF | P |
| --- | --- | --- | --- |
| Altitude | 2.03 | 1, 551.8 | 0.1552 |
| Plant origin | 12.97 | 1, 75.5 | **0.0006** |
| LDMC | 12.23 | 1, 16.9 | **0.0028** |
| SLA | 7.63 | 1, 22.7 | **0.0112** |
| Height | 2.22 | 1, 162.4 | 0.1382 |
| Altitude x Plant origin | 36.76 | 1, 1556.6 | **< $10^{-8}$** |
| Altitude x LDMC | 8.39 | 1, 665.4 | **0.0039** |
| Altitude x SLA | 10.40 | 1, 1576.8 | **0.0013** |
| Altitude x Height | 0.90 | 1, 1626.1 | 0.3430 |
| Plant origin x LDMC | 0.64 | 1, 19.5 | 0.4349 |
| Plant origin x SLA | 2.65 | 1, 26.1 | 0.1156 |
| Plant origin x Height | 11.80 | 1, 219.5 | **0.0007** |
| Altitude x Plant origin x LDMC | 0.06 | 1, 999.8 | 0.8126 |
| Altitude x Plant origin x SLA | 25.57 | 1, 1659.2 | **< $10^{-6}$** |
| Altitude x Plant origin x Height | 5.82 | 1, 1605.3 | **0.0160** |

We also used linear mixed effect models (LMM) to compare the values of individual traits between plants from the lowland and the highland and between potted and naturally growing plants. We fitted LMMs with individual traits as response variables, plant origin, altitude, and their interaction as predictors, and plant species, site, and cluster nested within site as random factors.

We evaluated statistical significance of fixed effects using F-tests implemented in the R package *lmerTest* (Kuznetsova et al. 2017). To test differences among plant species (a random effect), we compared LMMs with and without plant species using a likelihood ratio test.

## Results

Herbivory damage was driven by the interaction between the plant origin (potted lowland plants vs. plants naturally growing at both elevations) and the altitude (LMM, interaction plant origin x. altitude: F = 36.76, DF = 1, 1556.6, P <10^-8^; Table 3). While herbivory damage was reduced in the highland compared to the lowland in potted plants originating in the lowland, it was independent of the elevation in plants naturally growing at both elevations (Figs. 2 and 3).

**Figure 2.**
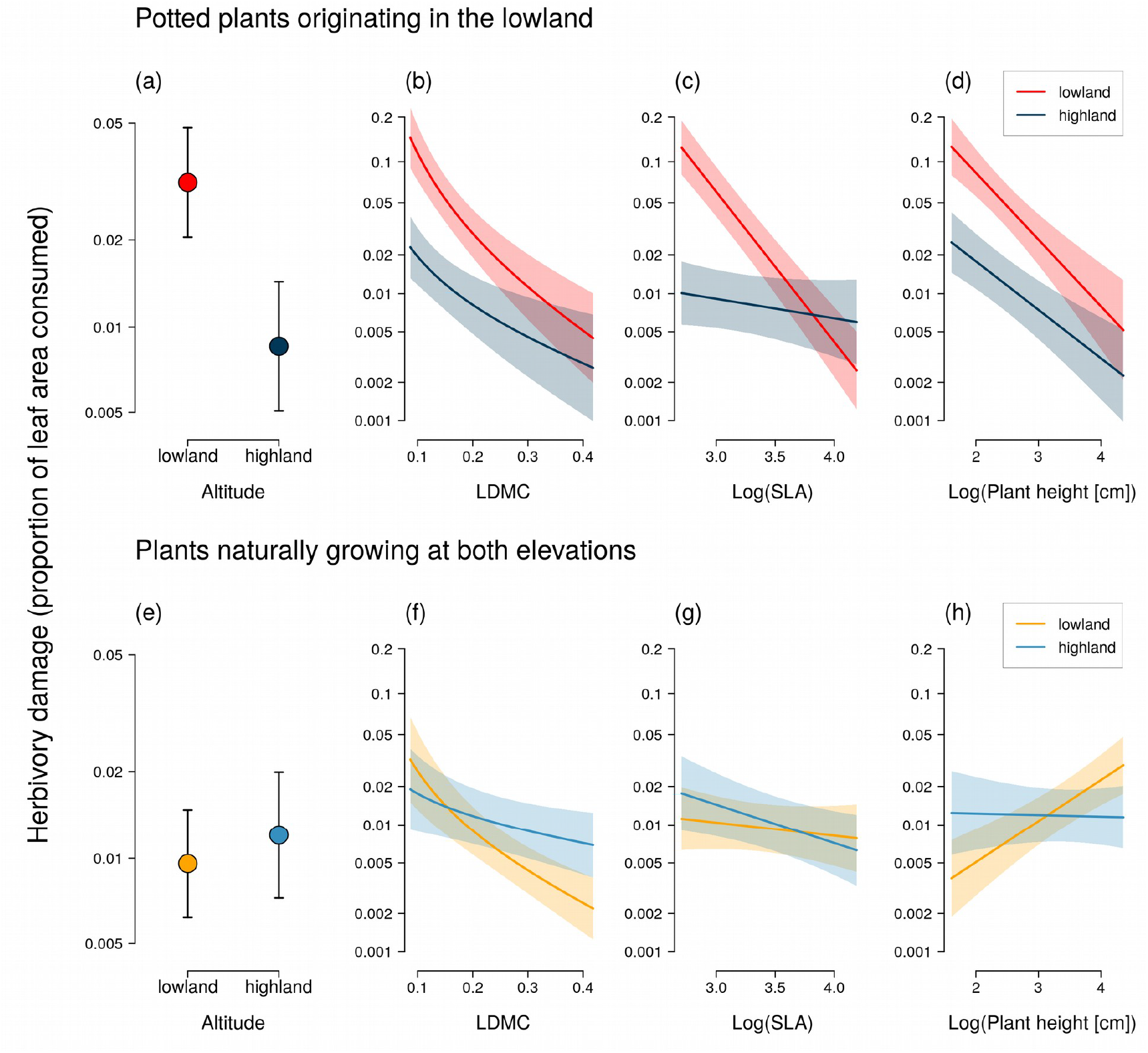
Herbivory damage at different altitudes was affected by plant origin and by functional traits. Potted plants originating in the lowland had lower herbivory damage (the proportion of leaf area consumed by herbivores) in the highland (a), while herbivory damage of plants naturally growing at both altitudes did not differ between the lowland and the highland sites (e). The effect of LDMC (b and f), SLA (c and g), and plant height (d and h) on herbivory damage in the lowland and the highland varied depending on the plant origin (potted lowland plants vs. plants naturally growing at both elevations). The plot shows estimates (mean ± standard error) from a generalised liner mixed effects model specified in Table 3, which included species, site, and cluster nested within site as random effects. The y-axis is displayed using the logit scale.

**Figure 3.**
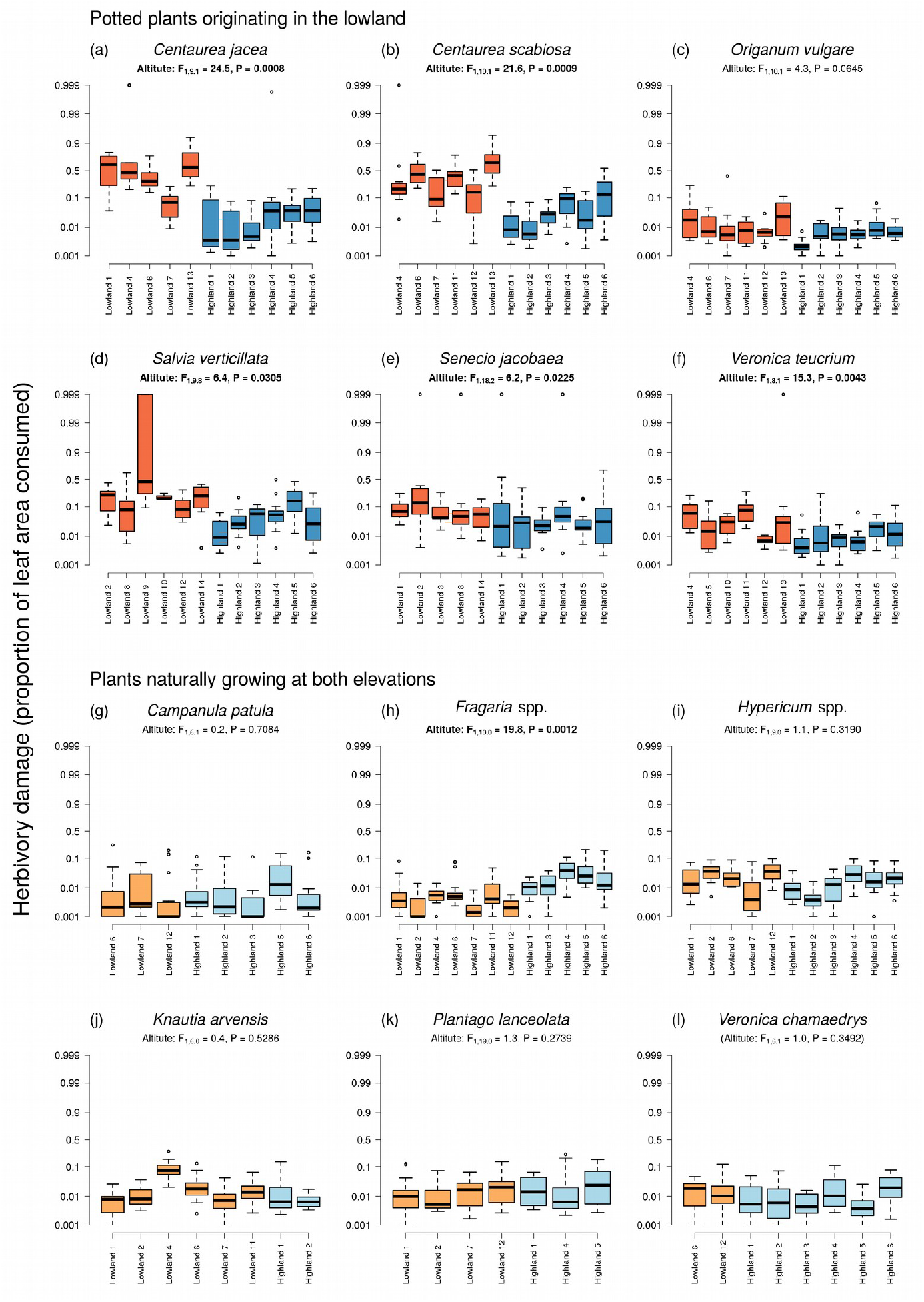
Herbivory damage of individual species at individual sites. Box plots show the values of herbivory damage (the proportion of leaf area consumed by herbivores) of individual species at the lowland (orange) and highland (blue) sites. Data for potted plants originating in the lowland are shown in the first and second row (a – f) and data for plants naturally growing at both elevations are shown in the third and fourth row (g – l). Results of linear mixed effects models comparing the herbivory damage in the lowland and the highland separately for each species are also provided (site and cluster nested within site used as random effects). The y-axis is displayed using the logit scale.

In addition, herbivory damage was also affected by plant height, SLA, and LDMC. We detected statistically significant interactions of plant height and SLA with plant origin and altitude (Table 3). However, herbivory damage mostly decreased with increasing SLA, increasing LDMC, and increasing plant height, although with differences in the slope between potted and naturally growing plants and between plants in the lowland and the highland (Fig. 2, Table 3). Mean herbivory damage differed among plant species (comparison of LMMs with and without species as a random effect: χ^2^ = 355.31, P <10^-15^, Fig. 3).

Measured plant traits varied among species, according to comparisons of LMMs with and without species as a random effect: height (χ^2^ = 2045.2, P <10^-15^), SLA (χ^2^ = 346.79, P <10^-15^), and LDMC (χ^2^ = 911.64, P <10^-15^). Both naturally growing and potted plants collected in the highland were on average taller by 4.52 cm (F = 5.21, DF = 1, 14.728, P = 0.0377), had SLA higher by 3.85 (F = 7.60, DF = 1,15.17, P = 0.0146), and LDMC lower by 0.022 (F = 7.23, DF = 1,15.027, P = 0.0168) compared to plants of the same species sampled in the lowland (Fig. 4). Potted plants were on average shorter by 14.51 cm compared to the naturally growing plants we sampled (F = 20.07, DF = 1, 10.01, P = 0.0012), had similar SLA (F = 0.41, DF = 1, 9.98, P = 0.5346), and LDMC lower by 0.107 (F = 11.74, DF = 1, 10.01, P = 0.0065).

**Figure 4.**
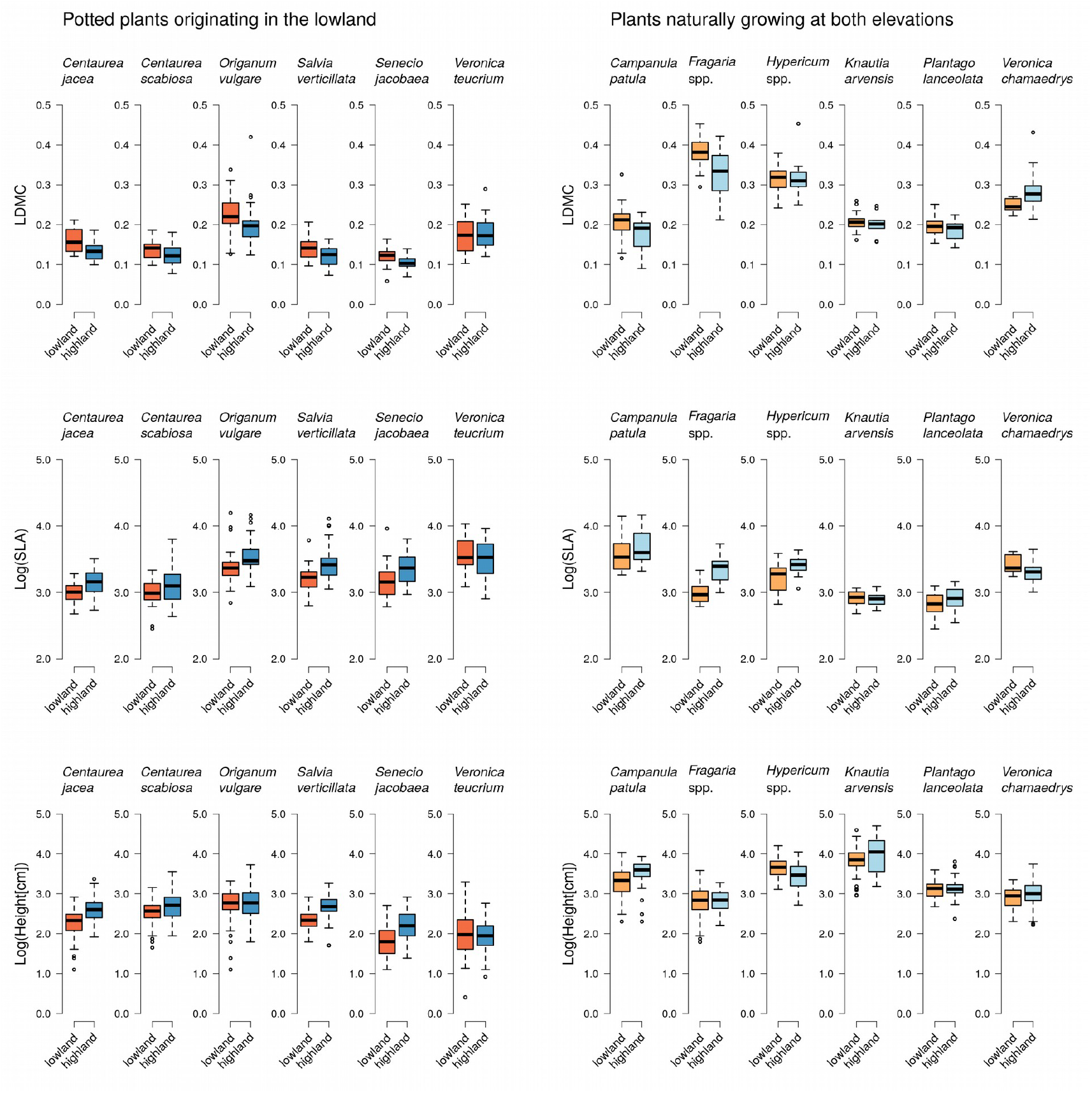
The values of functional traits measured in the lowland and the highland in individual plant species. The boxplots show the values of LDMC, SLA, and height of individual plants at all lowland (orange) and highland (blue) sites. Data for potted plants originating in the lowland are shown in the left column and data for plants naturally growing at both elevations are shown in the right column.

## Discussion

### Lowland plants had reduced herbivory damage when moved to higher altitudes

The results of our field experiment show that several thermophilous plant species originally growing in the lowland were significantly less damaged by invertebrate herbivores in the highland, based on the measurement of herbivory damage in individuals grown in pots. These results hold in comparison to control measurements of herbivory damage in plants naturally growing in both the lowland and the highland which revealed no significant difference in natural herbivory between the two altitudes.Our results thus provide support for the enemy release hypothesis (Costan et al. 2022; Kambo & Kotanen 2014) in a new context, specifically as a potential mechanism facilitating the uphill shift of plants induced by climate change.

The enemy release hypothesis was previously confirmed in exotic species, such as *Impatiens glandulifera* (Hejda & Pyšek 2006) and *Heracleum mantegazzianum* (Pyšek 1991) in Europe, which suffer decreased level of herbivory after they spread to new regions. Two main mechanisms of enemy release have been documented. First, a plant can escape its specialised herbivores which occur in its native range but not in the invasive range (Keane & Crawley 2002; Costan et al. 2022). It can still be consumed by generalised herbivores, but the overall herbivory damage is reduced because of the absence of specialised herbivores. The second mechanism, also called the novel weapon hypothesis, is that the exotic plant may contain defence compounds not present in plants growing in the invaded community, which means that local herbivores are evolutionarily naive to these defence compounds and cannot consume the exotic plant (Cappuccino & Carpenter 2005; Enge et al. 2012). This mechanism can reduce damage caused also by generalised herbivores.

Both mechanisms can explain the observations that plant species which are phylogenetically more distant from other native species in the community have a higher chance to become invasive (Omer et al. 2022). For this reason, it was previously unclear whether plants shifting their range over short distances, such as plants shifting to higher altitudes (Walther et al. 2005; Lenoir et al. 2008; Steinbauer et al. 2018), may also benefit from enemy release. The short geographic distance travelled by the plants mean that they are likely to encounter herbivores feeding on related species and thus preadapted to attack the new plants. However, our results clearly demonstrate that plants can benefit from reduced herbivory when they move to higher altitudes in the same region. The exact mechanism is not clear because we did not measure the abundance and community composition of herbivores at the two elevations. At the same time, the abundance of herbivores as such does not provide much information about the herbivore pressure on individual plant species without detailed knowledge of food preferences of individual species of herbivores. For more mechanistic insights, we would have to know the community food webs. Therefore we assume that the differences in herbivory damage between elevations were driven by differences in the composition of the herbivore communities in the highland compared to the lowland, such as lower abundance of slugs (Scheidel & Bruelheide 2001), which were apparently the main herbivores of *Centaurea jacea* and *C. scabiosa* in the lowland based on the type of leaf damage and slime trails we observed on the plants.

### No difference in herbivory damage of species naturally growing across the entire range of altitude

In comparison to lowland plants in the pots, the naturally growing plant species did not show significant differences in herbivory damage between lowland and highland sites. The only exception was *Fragaria* spp., but we do not have an explanation for this deviation. The naturally growing species we sampled are all common in grassland habitats and occur across the entire climatic gradient from the warmest lowlands to the highest mountain ranges in the Czech Republic (Chytrý et al. 2021). We can thus be confident that the populations we sampled occur in our study area for a long time and can be attacked by generalised herbivores as well as by coevolved specialists (Futuyma & Agrawal 2009; Maron et al. 2019). The six plant species/genera come from several different families (*Rosaceae, Campanulaceae, Plantaginaceae, Dipsacaceae, Hypericaceae*), which vary in defence mechanisms, and the comparison of herbivory damage of these plants between the lowland and the highland can thus provide a relatively robust estimate of the difference in the overall abundance and activity of herbivorous invertebrates between the two altitudes (Scheidel et al. 2003), with the caveat that we observed minor changes in functional traits between the lowland and the highland populations, which may affect their palatability. On the other hand, we observed the same changes in the potted plants, so this is unlikely to bias pour results.

A number of studies conducted in the temperate region reported the decrease of herbivory damage of naturally growing plants with increasing altitude (Reynolds & Crossley 1997; Scheidel et al. 2003; Pellisier et al. 2014, Galman et al. 2018), although this pattern is not universal and some studies reported no change of herbivory with altitude, increasing herbivory with altitude, or a hump-shaped relationship with a mid-elevation peak of herbivory damage (Scheidel & Bruelheide 2001; Scheidel et al. 2003; Moreira et al. 2017). It is assumed that when herbivory decreases at higher altitudes it is because of harsh environmental conditions, in particular lower temperature, which limits the abundance and activity of generalist herbivorous invertebrates (Scheidel & Bruelheide 2001; Scheidel et al. 2003; Moreira et al. 2017).

In our study, we compared only two elevations separated by ca. 400 m of altitude, because our aim was to experimentally test the effect of a realistic altitudinal shift of plant distribution on herbivory, rather than to provide a detailed description of changes of herbivory damage along an altitudinal gradient. Hence, it is likely that the lack of differences in herbivory damage of naturally growing plants we observed between the lowland and highland sites was caused by the relatively small difference in altitude leading to similar total abundance of herbivores.

### The effect of plant traits on herbivory damage

Overall, plant species differed in herbivory damage and all three measured traits, i.e., height, LDMC, and SLA, which reflects their different evolutionary histories translating into specific anti-herbivore strategies (Carmona et al. 2011; War et al. 2018). Differences in functional traits, particularly LDMC, are known to affect leaf palatability to herbivores (Schädler et al. 2003; Descombes et al. 2017) and variation of herbivory damage across different plant species (Schädler et al. 2003; Těšitel et al. 2021; Hrubá et al. 2022).

In our case, both potted and naturally growing plants collected in highlands were on average taller, had lower LDMC, and naturally growing plants also had higher SLA compared to individuals in the lowland. The fact that functional traits responded to altitude the same way in potted and naturally growing plants also supports our use of naturally growing plants as a control for the comparison of herbivory damage.

These general trends in functional traits indicate higher palatability of the highland plants in accordance with the findings of Descombes et al. (2017), who reported a decrease of LDMC and an increase in experimentally measured leaf palatability with the increasing altitude in the Swiss Alps, which may be related to lower abundance of herbivores in mountain habitats (Scheidel & Bruelheide 2001; Scheidel et al. 2003; Moreira et al. 2017). Plant growth and functional traits may be affected interactively by water and nutrient availability (Cossani et al. 2012; Akter & Klečka 2022), which we did not measure. However, potted plants were planted in a standardised soil mixture, which suggests that higher humidity in the highland habitats was likely driving these trait differences. Potted plants were on average shorter and had slightly lower LDMC than naturally growing plants, both of which simply follows from the fact that potted plants were younger, grown from seeds sown in the greenhouse ca. 3 months before the start of the experiment.

Herbivory rate was consistently negatively related to LDMC in both habitats and in both naturally growing and potted plants, which corresponds to previous findings (Schädler et al. 2003; Těšitel et al. 2021; Hrubá et al. 2022). Leaves with lower LDMC are usually more palatable for herbivores (Pálková & Lepš 2008; Descombes et al. 2017), likely because they are easier to chew. However, some herbivores may prefer tougher leaves with higher LDMC, as reported in locusts by Dostálek et al. (2020). In contrast to LDMC, the effect of plant height and SLA on herbivory damage was less consistent and was pronounced mostly in potted plants in the lowland, where a negative relationship between herbivory damage and both plant height and SLA was apparent. Potted plants were on average shorter than naturally growing plants, which makes them more susceptible to flightless herbivores such as slugs and snails which are likely more abundant in the lowland. These flightless invertebrates can consume whole seedlings but are less capable of reaching taller plants (Hahn et al. 2011), which are thus exposed to a restricted set of invertebrate herbivores.

Similarly to our results, Poorter et al. (2004) and Dostálek et al. (2020) recorded that herbivory damage decreased with increasing SLA. However, in our case, the negative effect of SLA was driven mostly by potted plants at the lowland sites, while the relationship between herbivore damage and SLA in naturally growing plants was weak or absent. This is consistent with the fact that SLA did not significantly influence herbivory damage in previous studies by Whitfeld et al. (2012) and Hrubá et al. (2022). On the other hand, multiple studies reported a positive relationship between SLA and herbivory damage (Dirzo 1980; Kozlov et al. 2015; Reese et al. 2016). Plants with higher SLA are also more attractive to slugs (Moshgani et al. 2017). Both SLA and LDMC serve as proxies for leaf toughness, which is functionally important for chewing herbivores. However, nutrient content and specific secondary metabolites also affect herbivory, but their effect is taxon-specific (Cornell & Hawkins 2003; Kelly & Bowers 2016). In our case, we measured only the total herbivore damage without identifying the herbivores feeding on the plants, so measuring additional traits, such as leaf chemistry, in more detail would not yield more insight into the interspecific differences in herbivory damage.

### Conclusions

Our main hypothesis, i.e., that lowland plant species would be less damaged by herbivores when they move to altitudes above their current upper altitudinal limit, was supported by the results of our experiment. Our data also show that plants with lower LDMC were less damaged by herbivores, while the effect of SLA and plant height was less pronounced and variable across altitudes. Overall, our results suggest that altitudinal shifts in the distribution of plant species may be facilitated by a decrease of herbivory damage in the newly colonized environment, as predicted by the enemy release hypothesis. More mechanistic insights could be obtained by future studies focusing on comparisons of species known to recently colonise mountain habitats with their native relatives and by studies taking the food web approach which includes information about food preferences of individual herbivore species. While reduced herbivory may facilitate the spread of plant species from lower altitudes to mountain habitats, long-term changes of mountain plant communities will also depend on interactions of plants with herbivores shifting to higher altitudes (Descombes et al. 2020). Improved understanding of these processes will be necessary to evaluate the impact of altitudinal shifts of species distribution on biodiversity of mountain habitats.

## Acknowledgements

We would like to thank Barbora Hazuková, Daniela Kameníčková, Lydie Plačková and Daniel Jackwerth for their help with experimental work in the field and with processing of leaf samples in the lab. We are grateful to the management of the Blanský Les Protected Landscape Area, Šumava Protected Landscape Area and all landowners for their cooperation and permission to conduct the study. We would also like to thank Ondřej Mudrák for his comments on the experimental design and on the previous version of the manuscript.

## Data availability

All data are available in Figshare: https://doi.org/10.6084/m9.figshare.22159244.

